# Inhibition of arenavirus entry and replication by the cell-intrinsic restriction factor ZMPSTE24 is enhanced by IFITM antiviral activity

**DOI:** 10.1101/2021.04.12.439453

**Authors:** Robert J Stott, Toshana L Foster

## Abstract

In the absence of effective vaccines and treatments, annual outbreaks of severe human haemorrhagic fever caused by arenaviruses, such as Lassa virus, continue to pose a significant human health threat. Understanding the balance of cellular factors that inhibit or promote arenavirus infection may have important implications for the development of effective antiviral strategies. Here, we identified the cell-intrinsic zinc transmembrane metalloprotease, ZMPSTE24, as a restriction factor against arenaviruses. Notably, CRISPR-Cas9-mediated knockout of ZMPSTE24 in human alveolar epithelial A549 cells increased arenavirus glycoprotein-mediated viral entry in pseudoparticle assays and live virus infection models. As a barrier to viral entry and replication, ZMPSTE24 may act as a downstream effector of interferon-induced transmembrane protein (IFITM) antiviral function; though through a yet poorly understood mechanism. Overexpression of IFITM1, IFITM2 and IFITM3 proteins did not restrict the entry of pseudoparticles carrying arenavirus envelope glycoproteins and live virus infection, yet depletion of IFITM protein expression enhanced virus entry and replication. Furthermore, gain-of-function studies revealed that IFITMs augment the antiviral activity of ZMPSTE24 against arenaviruses, suggesting a cooperative effect of viral restriction. We show that ZMPSTE24 and IFITMs affect the kinetics of cellular endocytosis, suggesting that perturbation of membrane structure and stability is likely the mechanism of ZMPSTE24-mediated restriction and cooperative ZMPSTE24-IFITM antiviral activity. Collectively, our findings define the role of ZMPSTE24 host restriction activity in the early stages of arenavirus infection. Moreover, we provide insight into the importance of cellular membrane integrity for productive fusion of arenaviruses and highlight a novel avenue for therapeutic development.

**Author Summary:** Increased human travel, virus genome evolution and expansion of the host rodent reservoir outside of endemic areas has contributed to increasing cases of the highly fatal arenaviral haemorrhagic disease, Lassa fever in Western Africa. These annual seasonal outbreaks present a serious global public health and socioeconomic burden, particularly in the absence of approved vaccines and antiviral countermeasures. Development of novel and effective therapeutic strategies against arenavirus infection is reliant on a better understanding of the molecular mechanisms of key host–virus interactions that antagonise or potentiate disease pathogenesis. We demonstrate the inhibition of arenavirus infection by the antiviral restriction factor ZMPSTE24 and describe a cooperative action with the innate immunity-stimulated family of interferon-induced transmembrane proteins (IFITMs). This work adds to our understanding of the mechanism of ZMPSTE24 and IFITM-mediated restriction of enveloped viruses and importantly suggests that these proteins may play a significant role in the pathogenesis of arenavirus infections.

## Introduction

Viral haemorrhagic fevers (VHF) caused by mammarenaviruses pose a serious public health burden in endemic regions. Based on their phylogenetic and antigenic properties and geographical distribution, mammarenaviruses are classified into the Old World (OW) virus complex endemic to Western Africa and the New World (NW) virus complex found in South America (1, 2). Junín virus (JUNV), the causative agent of Argentinian haemorrhagic fever (AHF) is included within the NW group along with the other human pathogens Machupo (MACV) and Chapare (CHAPV) viruses (3, 4). The OW group includes the clinically significant, prototypic arenavirus lymphocytic choriomeningitis virus (LCMV) which is distributed worldwide, and the pathogenic Lujo virus (LUJV) associated with an outbreak of VHF in South Africa and Zambia (5). The most prevalent mammarenavirus pathogen, OW Lassa virus (LASV) is the causative agent of human Lassa Fever (LF) and is associated with significant mortality and morbidity in affected West African countries. Increased human interactions with the rodent host reservoir, mainly *Mastomys natalensis*, have led to the emergence and re-emergence of LF disease (6–9). Particularly Nigeria reports continued outbreaks of LF, resulting in tens of thousands of cases and lead to case fatality rates as high as 30% (10, 11). Recent expansion of LASV outside of endemic regions has further highlighted the immense risk to human health in the absence of effective and approved countermeasures (12–15). A distinct characteristic of OW and NW mammarenaviruses is that both groups contain human pathogenic and non-pathogenic strains (1). For example, OW Mopeia virus (MOPV) is known to be non-pathogenic to humans, yet it is phylogenetically closely related to LASV and has been shown to induce protective immunity against LASV in a non-human primate model (16–18). In the case of LASV, infections can be asymptomatic or result in illnesses ranging from mild flu-like syndromes to severe and highly fatal haemorrhagic zoonoses (19). Thus, unravelling the immune responses and key virus-host interactions that influence this variation in disease severity is critical.

The innate antiviral response provides an early line of defense against infection through the activity of cell-intrinsic proteins and the induction of type I interferon (IFN1)–stimulated genes (ISGs) that encode virus restriction factors (20, 21). These innate restriction factors block specific steps of the viral life cycle, including fusion of the viral membrane of enveloped viruses, a critical determinant for establishing infection (21–24). Fusion is mediated by the viral fusion glycoproteins that usually undergo conformational changes upon receptor binding and/or induction by acidic pH in endosomal compartments of the cell (25–28). The viral and cellular membrane leaflets merge to form a hemifusion diaphragm. Lipid mixing between the two leaflets then leads to the formation of a fusion pore, through which viral content is released into the cytoplasm to initiate the replication process (29, 30).

Several restriction factors potently inhibit the infection of diverse enveloped viruses (31, 32). Amongst these factors are the interferon-induced transmembrane proteins (IFITMs) and the zinc metalloprotease, ZMPSTE24 (33, 34). As part of the robust IFN-mediated innate immune response to viral infection, the IFITMs 1, 2 and 3 display broad-range antiviral activity against a plethora of enveloped viruses including orthomyxoviruses (influenza A virus, IAV), retroviruses (human immune deficiency virus-1, HIV-1), coronaviruses (severe acute respiratory syndrome (SARS) coronaviruses SARS-COV-1 and SARS-CoV-2), flaviviruses (Dengue) and filoviruses (Ebola) (35–39). IFITM1 localises predominantly to the plasma membrane, whilst IFITMs 2 and 3 localise to early and late endosomal and lysosomal membranes (21, 40). Current mechanisms of virus restriction are not clearly understood but existing explanations suggest that IFITMs inhibit viral fusion through a proximity-based mechanism (36, 39). This is proposed to require the presence of IFITMs at the site of viral fusion where they inhibit the formation of the fusion pore by trapping this process at the hemifusion stage. IFITM homo-oligomerisation is thought to directly modify the structure, rigidity and curvature of target membranes, thus leading to a block in virus-host fusion (36, 41). Indirect mechanisms have also been suggested, such as through alteration of membrane cholesterol composition or through endosomal association with other membrane proteins, such as ZMPSTE24 (34).

ZMPSTE24, a seven-pass transmembrane protein, has recently been shown to be an intrinsic restriction factor against a number of enveloped viruses, including IAV, Ebola, vaccinia and Zika (33). ZMPSTE24 is constitutively expressed and localises to the inner nuclear membrane, and to multiple intracellular endocytic membrane compartments. The conserved enzymatic activity of ZMPSTE24 plays a crucial role in the maturation of the nuclear scaffold protein lamin A which is critical for nuclear structure, shape and function (42). Recent studies suggest an indirect mechanism of ZMPSTE24 inhibition of enveloped virus fusion, independent of this enzymatic activity, through alteration of membrane structure or endosomal trafficking, similar to the IFITM proteins (33). ZMPSTE24 has been identified as a protein-interaction partner of IFITMs and recruitment of ZMPSTE24 as a downstream effector of IFITM antiviral activity has been suggested to drive the alteration in membrane properties that are less conducive to viral fusion (34). However, the exact mechanism of this proposed cooperative restriction is currently unknown.

Different viruses can initiate fusion in distinctive endosomal and lysosomal compartments and thus can display differential sensitivity to restriction factor expression. Arenavirus entry into target cells is mediated by the glycoprotein spike complex GP, consisting of subunits GP1, GP2 and the stable signal peptide (SSP). GP1 binds to entry receptors at the plasma membrane and effective fusion is driven by stabilisation of receptor-GP complexes by GP2 and SSP (43). During OW and NW virus co-divergence, cell receptor usage evolved from the ubiquitously expressed α-dystroglycan receptor (OW) to the transferrin 1 receptor (NW), with the exception of OW LUJV which uses neuropilin-2 (NRP-2) as the entry receptor (28, 44). Virus entry occurs either via clathrin-dependent (NW viruses) or independent receptor-mediated (OW viruses) endocytosis followed by low-pH induced conformational changes in GP structure that lead to productive fusion (45). For OW LASV, low pH triggers a switch from the α-dystroglycan receptor to the lysosome-associated membrane protein 1 (LAMP1), whilst for OW LUJV a pH-induced switch to the tetraspanin CD63 mediates fusion with cellular membranes (26).

To date, there is limited information about how arenavirus infections may be antagonised by host restriction factors (32, 46, 47), and also the putative role of ZMPSTE24 in arenavirus biology has not yet been explored. Given that ZMPSTE24 is an important innate defence factor against a number of pathogenic viruses, it is conceivable that ZMPSTE24 may affect the viral endocytic entry process of arenaviruses. In this study, we examined whether ZMPSTE24 is involved in arenavirus entry restriction. Using complementary arenavirus GP-pseudoparticle (GPpp) and live MOPV infection assays, we demonstrated that ZMPSTE24 restricts the entry and replication of arenaviruses. In agreement with previous reports, we found that arenavirus entry was resistant to IFITM protein overexpression (39, 48). However, siRNA knockdown and CRISPR-Cas9 knockout of IFITMs enhanced arenavirus entry and replication, contrary to our expectations. We found that IFITM3 overexpression in the presence of ZMPSTE24 augmented restriction of arenavirus entry and replication and showed that ZMPSTE24 and IFITM3 can alter cellular endocytosis rates possibly by impacting on the rigidity of cell membranes in an independent or cooperative manner. Collectively, our results provide strong support that arenaviruses utilise an endocytic pathway that is sensitive to ZMPSTE24 restriction and is enhanced upon recruitment of IFITM3.

## Results

### ZMPSTE24 impairs arenavirus GPpp infection

The broad-spectrum intrinsic restriction factor ZMPSTE24 blocks the endocytic entry of a number of enveloped viruses (33). We used arenavirus GP–pseudoparticles (GPpp) generated from a panel of OW (LCMV, LASV, LUJV, MOPV) and NW (JUNV, MACV, CHAPV) mammarenavirus representatives to examine the role of ZMPSTE24 restriction during arenavirus entry. Specifically, murine leukaemia virus (MLV) packaging a green fluorescent protein (GFP) reporter was pseudotyped with different arenavirus GP proteins. Using flow cytometry analysis, we first investigated the infectivity of these arenavirus GPpp in A549 human lung epithelial cells stably over-expressing FLAG-tagged ZMPSTE24 compared to vector control cells. Entry of both OW and NW arenavirus GPpp were similarly susceptible to restriction by overexpression of ZMPSTE24 (Fig 1A). To examine the importance of ZMPSTE24 metalloprotease activity in the restriction of arenavirus GPpp entry, we mutated histidine residue 335 within the essential, conserved HEXXH zinc metalloprotease catalytic motif and demonstrated that the H335A mutant also displayed comparable restriction of LCMVpp and LASVpp infection to wild-type ZMPSTE24 in A549 cells when compared with vector only controls, albeit to a slightly lesser extent (Fig 1B). Therefore, the combined data suggest that ZMPSTE24 impedes arenavirus infection but that this viral restriction activity is largely independent of its protease function.

**Fig 1.**
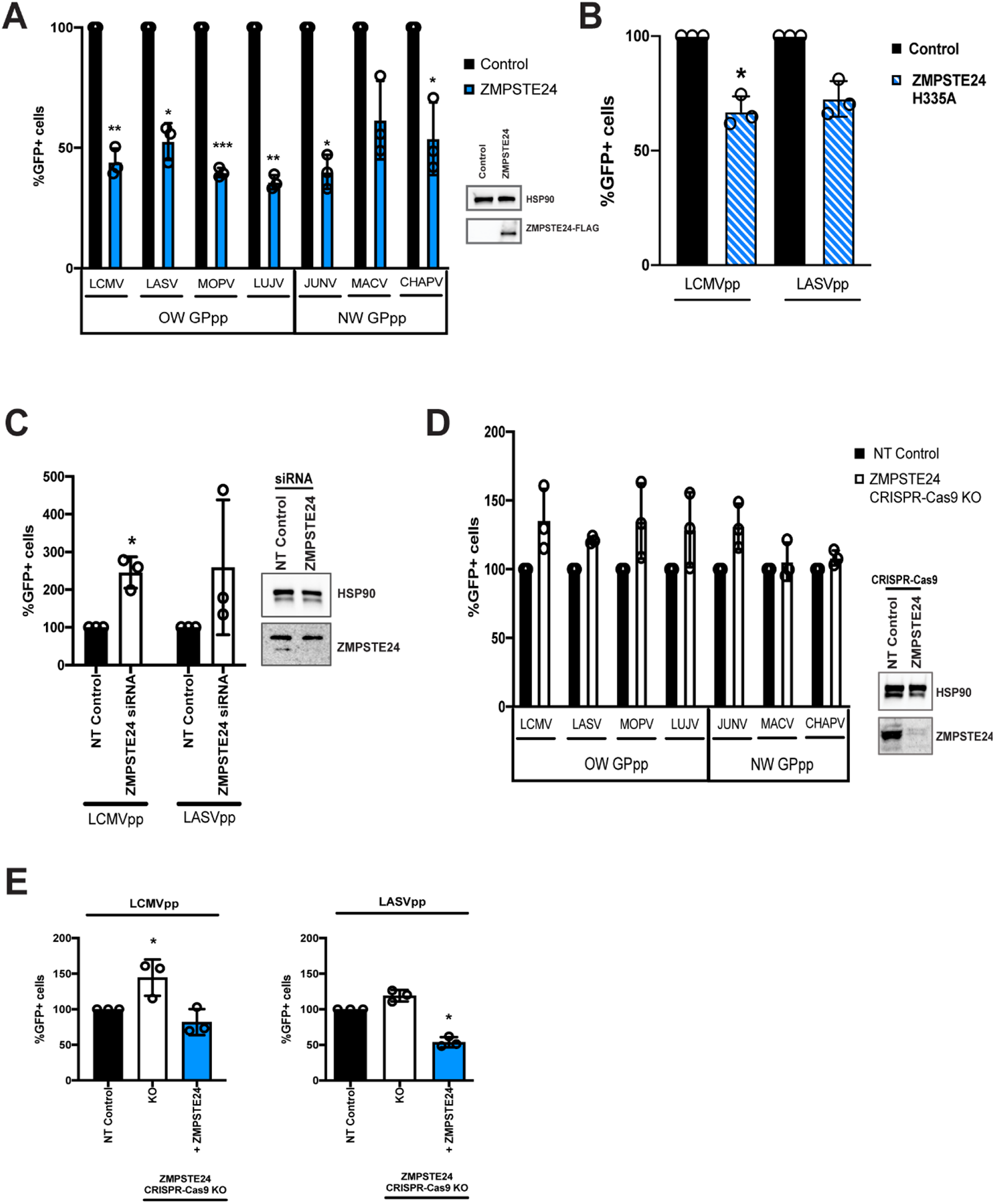
Entry of arenavirus GPpp is inhibited by ZMPSTE24. GFP-containing arenavirus GP-pseudoparticles (GPpp) generated from a panel of Old World (OW) and New World (NW) arenaviruses namely, LCMV, LASV, MOPV, LUJV, JUNV, MACV and CHAPV were used to infect: (A) A549 cells stably transduced with vector control or ZMPSTE24-FLAG (inset), (B) A549 cells stably expressing vector control or ZMPSTE24 with a H335A mutation, (C) A549 cells with ZMPSTE24 expression silenced by siRNA knockdown (inset, HSP90 served as a loading control), (D) CRISPR-Cas9 mediated knockout (KO) of ZMPSTE24 in A549 cells (inset) or (E) A549 cells stably transduced with non-targeting (NT) control, CRISPR-Cas9 KO of ZMPSTE24 or CRISPR-Cas9 KO of ZMPSTE24 in combination with ZMPSTE24-FLAG overexpression. Sensitivity to ZMPSTE24 restriction was measured by flow cytometry and is expressed as % GFP positive cells in relation to corresponding controls. Unpaired t-test *** p<0.001, ** p<0.01, *p<0.05. Data are expressed as mean ±SEM from samples produced in triplicate.

To corroborate the inhibitory effect of ZMPSTE24 on the entry of arenaviruses, we silenced the expression of ZMPSTE24 by performing siRNA knockdown (KD) studies in A549 cells. We found that LCMVpp and LASVpp infection was enhanced 2.5-fold in ZMPSTE24 KD cells compared to the scramble control siRNA (Fig 1C). To build on the siRNA knockdown data, given that the knockdown by western blot was apparently not complete, we depleted the expression of ZMPSTE24 in A549 cells using CRISPR-Cas9 lentiviral vectors. Cells were screened by western blotting to confirm the depletion of ZMPSTE24 expression in the knockout (KO) cells. Infection by most OW and NW arenavirus GPpp was markedly increased in the ZMPSTE24 knockout cells compared to unmodified control cells (Fig 1D). To further confirm the role of ZMPSTE24 as an intrinsic host factor against arenavirus GPpp entry, we modified the ZMPSTE24 KO A549 cells to stably express C-terminally FLAG-tagged ZMPSTE24 by retroviral transduction (Fig 1E). We observed less infection with LCMVpp and LASVpp in the ZMPSTE24-expressing cells compared to the ZMPSTE24 KO cells. Collectively, by using single-round GPpp infection assays, we have identified ZMPSTE24 as a host restriction factor against arenavirus entry.

### Arenavirus GPpp entry is resistant to IFITM restriction but depletion of IFITM expression enhances infection

It has been suggested that ZMPSTE24 is recruited to endocytic compartments by IFITM proteins, thereby blocking the endocytic entry of enveloped viruses (33, 34). We aimed to examine the role that IFITMs play in the restriction activity against arenaviruses. As IFITMs are induced by IFN stimulation, we first assessed the antiviral effects of exogenous IFN1 on the early stages of arenavirus infection in A549 cells. As measured by flow cytometry, single-round infectivity of GPpp across OW and NW strains was markedly reduced in cells incubated with 1000U/ml universal IFN1, suggesting an IFN-mediated inhibition of arenavirus entry (Fig 2A). It has previously been reported that arenaviruses are not susceptible to inhibition by the broadly acting IFN1-stimulated family of IFITM proteins (39, 48). To further validate this, we generated A549 cells stably expressing individual human IFITM1, IFITM2 and IFITM3 proteins at levels similar in magnitude to that induced by IFN1 treatment (Fig 2B). By contrast, the expression of endogenous ZMPSTE24 was not upregulated following treatment with IFN1. We analysed the infectivity of different arenavirus GPpp in these cells and found that all strains tested were resistant to antiviral IFITM activity (Fig 2C), which is in agreement with previous studies (39, 48).

**Fig 2.**
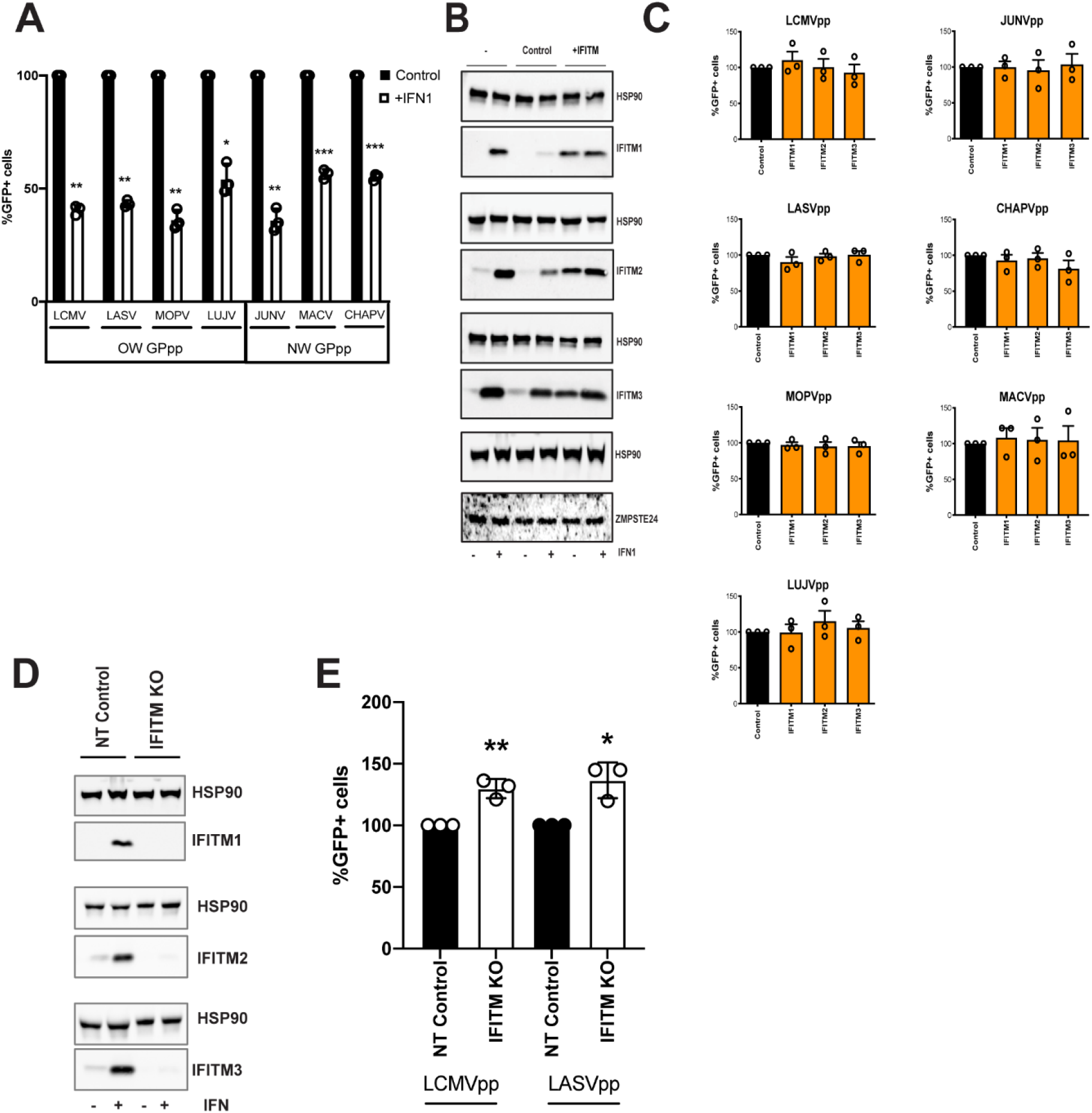
**Arenavirus GPpp are not susceptible to restriction by IFITM overexpression**. **(A)** A549 cells were infected with arenavirus GPpp for 48 h in the presence of 1000 U/mL type 1 IFN (IFN1) or control media and the percentage of infected cells compared to control was determined by flow cytometry. Unpaired t-test *** p<0.001, ** p<0.01, *p<0.05. **(B)** A549 cells were transduced with empty vector control pLHCX, IFITM1, IFITM2 or IFITM3. Expression of IFITMs and ZMPSTE24 was measured in the presence or absence of 1000 U/mL IFN1 by western blot analysis. HSP90 served as a loading control. **(C)** A549 cells stably transduced with IFITMs were infected with arenavirus GPpp and % infectivity measured by flow cytometry. **(D)** CRISPR-Cas9 mediated knockout (KO) of IFITMs in A549 cells was confirmed by western blot analysis in the presence or absence of 1000 U/mL IFN1. **(E)** A549 IFITM CRISPR-Cas9 KO cells were infected with LCMV or LASV GPpp for 48 h and the percentage of infected cells compared to control was determined by flow cytometry. Unpaired t-test *p<0.05. Data are expressed as mean ±SEM from experiments in triplicate.

To assess whether the depletion of endogenous IFITMs may alter the efficiency of arenavirus GPpp infection, we used CRISPR-Cas9 lentiviral vectors to target IFITM expression in A549 cells. As observed by western blotting, the IFITM3 guide RNA (gRNA) used resulted in effective depletion of the specific IFITM targeted but also resulted in the depletion of IFITM2 and IFITM1, given their high degree of homology, chromosomal positioning and close proximity (Fig 2D). Knockout of endogenous IFITM expression enhanced the infection mediated by LCMVpp and LASVpp, implying that IFITMs may contribute to the inhibition of arenavirus GP-mediated viral entry, possibly through co-factor interactions (Fig 2E).

### IFITMs contribute to the antiviral restriction of arenavirus entry by ZMPSTE24

To address the effects of IFITMs on ZMPSTE24 activity, we first assessed the comparative localisation of the host restriction factors. Immunofluorescence microscopy imaging showed that overexpressed HA-tagged ZMPSTE24 possesses a cytoplasmic distribution and predominantly localises to endosomal compartments in A549 cells. Further, we found that ZMPSTE24 co-localises with the early endosome marker, EEA1 and to a lesser extent with the late endosome marker Rab9, but does not localise to the lysosome marker LAMP1 (Fig S1). The localisation of IFITM proteins is thought to define the spectrum of viruses that these antiviral factors restrict (23). IFITM1 is mostly localised to the plasma membrane, whilst IFITMs 2 and 3, like ZMPSTE24, are localised to endosomal compartments due to the presence of a conserved endocytic localisation motif (21, 23). We further demonstrated by confocal microscopy imaging that endogenous ZMPSTE24 co-localises with all three HA-tagged IFITM proteins when overexpressed in A549 cells (Fig 3A). In the presence of IFITM1, ZMPSTE24 redistributed to the plasma membrane and IFITM1 was also found to have a disperse intracellular punctate distribution that overlapped with ZMPSTE24. This observation implies a cooperative function or interaction of the two proteins (Fig 3A). Considering these observations, we next corroborated that IFITMs and ZMPSTE24 interact (33, 34). A549 cells were transfected with C-terminally FLAG-tagged ZMPSTE24. Proteins were captured on beads coated with anti-IFITM2/3 antibody and analysed by western blot. IFITM proteins have the propensity to homo- and hetero-oligomerise and we found that ZMPSTE24-FLAG binds to the endogenous IFITM1, 2 and 3 proteins (Fig 3B) (41). Complementary to these co-immunoprecipitation data, we assessed the interaction between ZMPSTE24 and specifically, IFITM3, in live cells using a NanoLuc Binary Technology (NanoBiT)-based assay (Fig 3C). NanoLuc Luciferase is split into two complementary segments, 18kDa Large BiT (LgBiT) and 1.3kDa Small BiT (SmBiT); these possess low intrinsic affinity for each other. However, a bright luminescent signal is restored upon interaction of the binding partners to which they are fused. We engineered ZMPSTE24 and IFITM3 constructs tagged at either the N or C terminus with SmBiT and LgBiT fragments (Fig 3C). We transiently transfected HEK293T cells with these combinations of SmBiT/LgBiT ZMPSTE24 and IFITM3 constructs to screen for conformational interactions by detection of a luminescence signal. As IFITMs are known to oligomerize via the N-terminus, we included a co-transfection of N-terminal LgBiT-IFITM3 (N-L-IFITM3) and N-terminal SmBiT-IFITM3 (N-S-IFITM3) as indication of a positive interaction. Luminescence signal was also compared to the manufacturer’s PRKACA:PRKAR2A positive control pair and the negative control of the corresponding SmBiT partner fused with a HaloTag. Co-transfection of N-L-IFITM3 and N-S-IFITM3 produced a robust signal when normalised to the negative control, indicating the oligomerisation of IFITM proteins (Fig 3C). We found that C-terminal tagged ZMPSTE24 (ZMPSTE24-C-S) and IFITM3 (IFITM3-C-L) in combination produced a luminescent signal approximately 12-fold higher than that of the negative control pairs and approaching the signal of the N-L-IFITM3/N-S-IFITM3 combination (Fig 3C). Thus, these data are in keeping with previous findings that suggest the two host restriction factors may interact (33).

**Fig 3.**
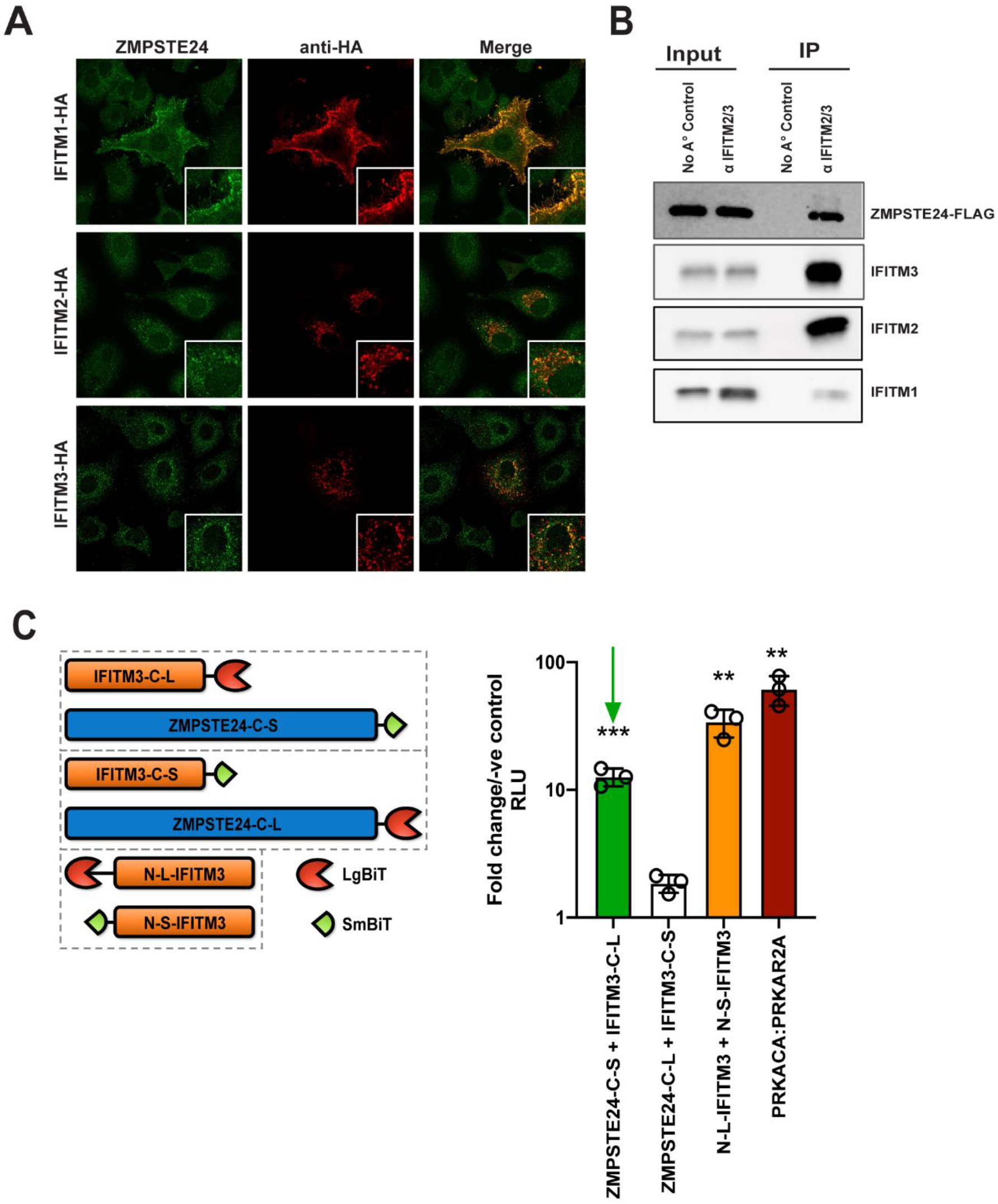
ZMPSTE24 colocalises with IFITM proteins, and interacts via a C-terminal interaction with IFITM3. **(A)** Confocal microscopy of A549 cells transiently transfected with HA-tagged IFITM proteins 1, 2 and 3. At 24 h post-transfection, cells were fixed and stained for endogenous ZMPSTE24 (green) and the IFITM protein of interest (red) and examined by confocal microscopy. Panels are of representative images. **(B)** A549 cells were transfected with ZMPSTE24-FLAG and 48 h later, cell lysates were immunoprecipitated with anti-IFITM2/3 monoclonal antibody. Cell lysates and immuprecipitates were analysed by SDS-PAGE and western blotting for ZMPSTE24-FLAG and IFITMs 1, 2 and 3. **(C)** IFITM3 or ZMPSTE24 were fused to a large (LgBiT) or small (SmBiT) subunit of NanoLuc luciferase and co-transfected into HEK293T cells. For each assay pair (hatched boxes), the LgBiT fused partner was also co-transfected with a HaloTag fused to SmBiT as a negative control. LgBiT-PRKACA and SmBiT-PRKAR2A were co-transfected as a positive control. **(D)** Luminescence was measured after addition of NanoGlo luciferase reagent and is expressed as relative luminescence units (RLU) relative to corresponding LgBiT and SmBiT-HaloTag negative control. Unpaired t-test *** p<0.001, ** p<0.01, * p<0.05. Data are expressed as mean ±SEM from experiments in triplicate. Green arrow indicates interaction between ZMPSTE24-C-SmBiT and IFITM3-C-LgBiT.

We next aimed to ascertain if IFITMs play a role in the ZMPSTE24-mediated restriction of arenavirus entry. We assessed the infectivity of LCMVpp and LASVpp in A549 cells with CRISPR-Cas9 KO of endogenous ZMPSTE24 and overexpressing either ZMPSTE24-FLAG or IFITM3, or both proteins together by retroviral transduction (Fig S2, Fig 4A). As anticipated, ZMPSTE24 knockout increased LCMVpp and LASVpp infection but this was abrogated in the presence of ZMPSTE24-FLAG. IFITM3 overexpression alone had little effect on GPpp infection but in combination with ZMPSTE24-FLAG, we observed a significant enhancement in the restriction of LCMVpp and LASVpp infection (Fig 4B). Interestingly, when we examined A549 cells overexpressing ZMPSTE24 and IFITM3 in combination, we found that ZMPSTE24 co-expression led to a redistribution of IFITM3 from the disparate endo-cytoplasmic localisation to a distinct endosomal localisation that overlaps with ZMPSTE24 expression (Fig 4B). Thus, we speculate that this redistribution of IFITM3 likely influences the observed enhancement in ZMPSTE24 restriction of arenavirus entry.

**Fig 4.**
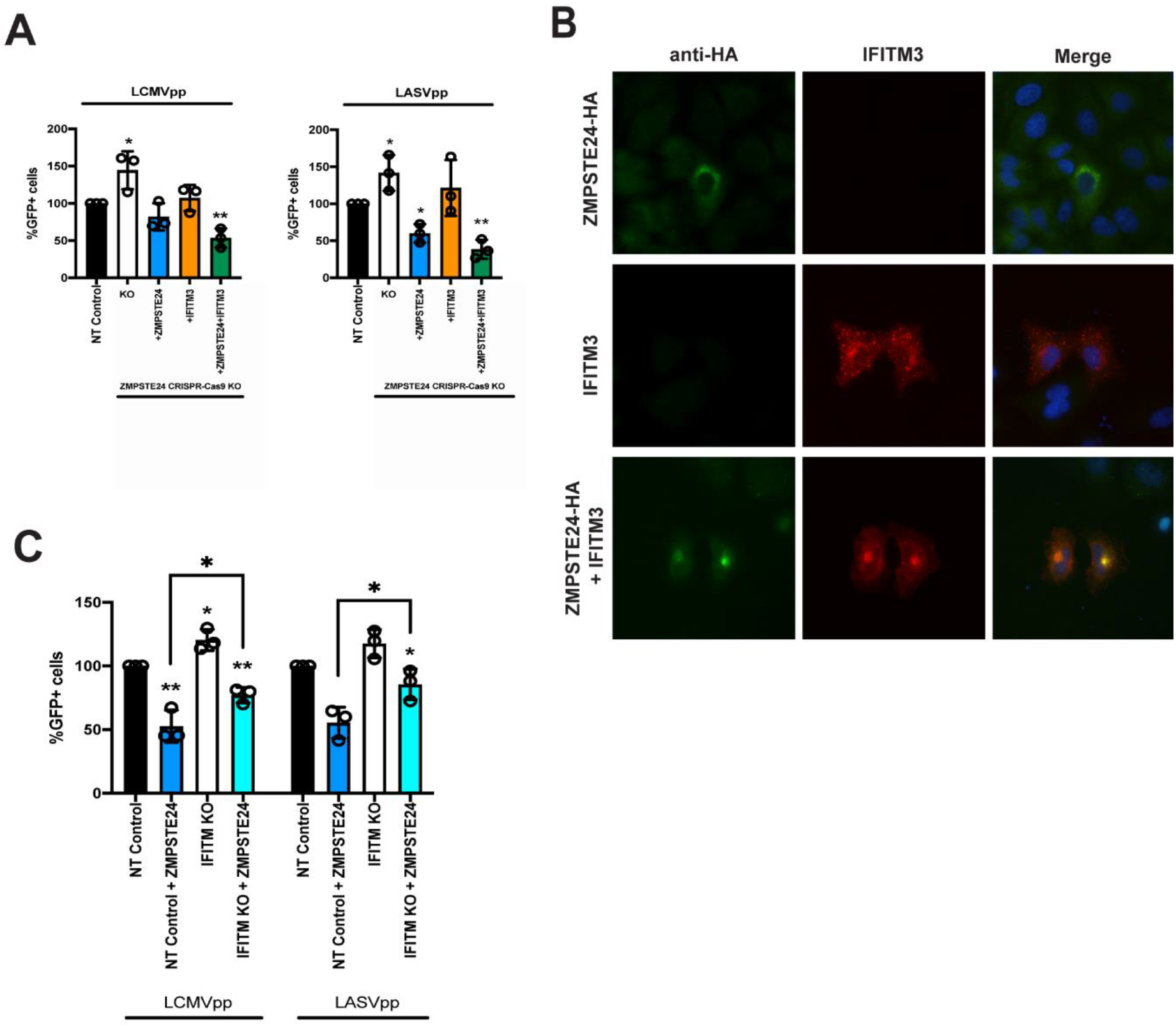
ZMPSTE24 and IFITM3 co-operatively restrict arenavirus GPpp infection. **(A)** A549 CRISPR-Cas9 stable knock out cell lines were generated for non-targeting (NT) control or ZMPSTE24 and then transduced for stable overexpression of ZMPSTE24-FLAG or IFITM3 or both. Cells were infected with LCMVpp or LASVpp for 48 h and the percentage of infected cells compared to control was determined by flow cytometry. Unpaired t-test ** p<0.01, *p<0.05. **(B)** A549 cells transiently transfected with HA-tagged ZMPSTE24 (green) and IFITM3 (red) were stained to assess the co-localisation of the two proteins 24 h post-transfection. Panels are of representative images. **(C)** A549 CRISPR-Cas9 NT control or IFITM KO cells ectopically expressing ZMPSTE24-FLAG were infected with LCMVpp or LASVpp. At 48 h post transfection the percentage of infected cells compared to control was determined by flow cytometry. Unpaired t-test ** p<0.01, *p<0.05. Data are expressed as mean ±SEM from samples produced in triplicate.

We next generated A549 lentiviral non-targeting (NT) control and CRISPR-Cas9 IFITM knockout cells and stably expressed ZMPSTE24-FLAG in these cells. Compared to NT control cells, we observed enhanced LCMVpp and LASVpp infection in IFITM KO cells (Fig 4C). Overexpression of ZMPSTE24 in NT control cells abrogated LCMVpp and LASVpp infection but in the absence of IFITM expression, the sensitivity of arenavirus entry to ZMPSTE24 restriction was decreased (Fig 4C). Taken together, our data show that ZMPSTE24 interacts with IFITM proteins and importantly suggest a cooperative impairment of arenavirus entry in which ZMPSTE24 appears to co-opt IFITM3 to facilitate restriction of arenavirus entry.

### Replication of OW MOPV is restricted by ZMPSTE24

We established that ZMPSTE24 can restrict the early stages of arenavirus infection through the use of GPpp infection assays and found that ZMPSTE24 interaction with IFITM proteins markedly increased the sensitivity of arenavirus-mediated entry to ZMPSTE24, suggesting a cooperative function. The replication of MOPV is highly sensitive to IFN1 inhibition, thus we next wanted to determine whether IFITM-mediated restriction contributed to this inhibition and to assess the impact of ZMPSTE24 on live virus replication (Fig 5A). Using lentiviral CRISR-Cas9 ZMPSTE24 gRNA and siRNAs against IFITMs 1, 2 and 3, we first knocked out or knocked down endogenous protein expression from A549 cells and challenged the cells with MOPV (Fig 5B, C). Levels of MOPV NP and L RNA in these cells were quantified by RT-qPCR at 72 hours post-infection. We found that ZMPSTE24 depletion markedly increased the level of viral replication observed. This was reflected in the 4-6-fold increase in MOPV NP and L RNA levels in the absence of ZMPSTE24 expression, indicating that in these cells ZMPSTE24 is likely a component of the induced antiviral state upon arenavirus infection (Fig 5B). Likewise, knockdown of IFITM protein expression, particularly of IFITMs 2 and 3 significantly enhanced MOPV NP and L RNA levels (Fig 5C).

**Fig 5.**
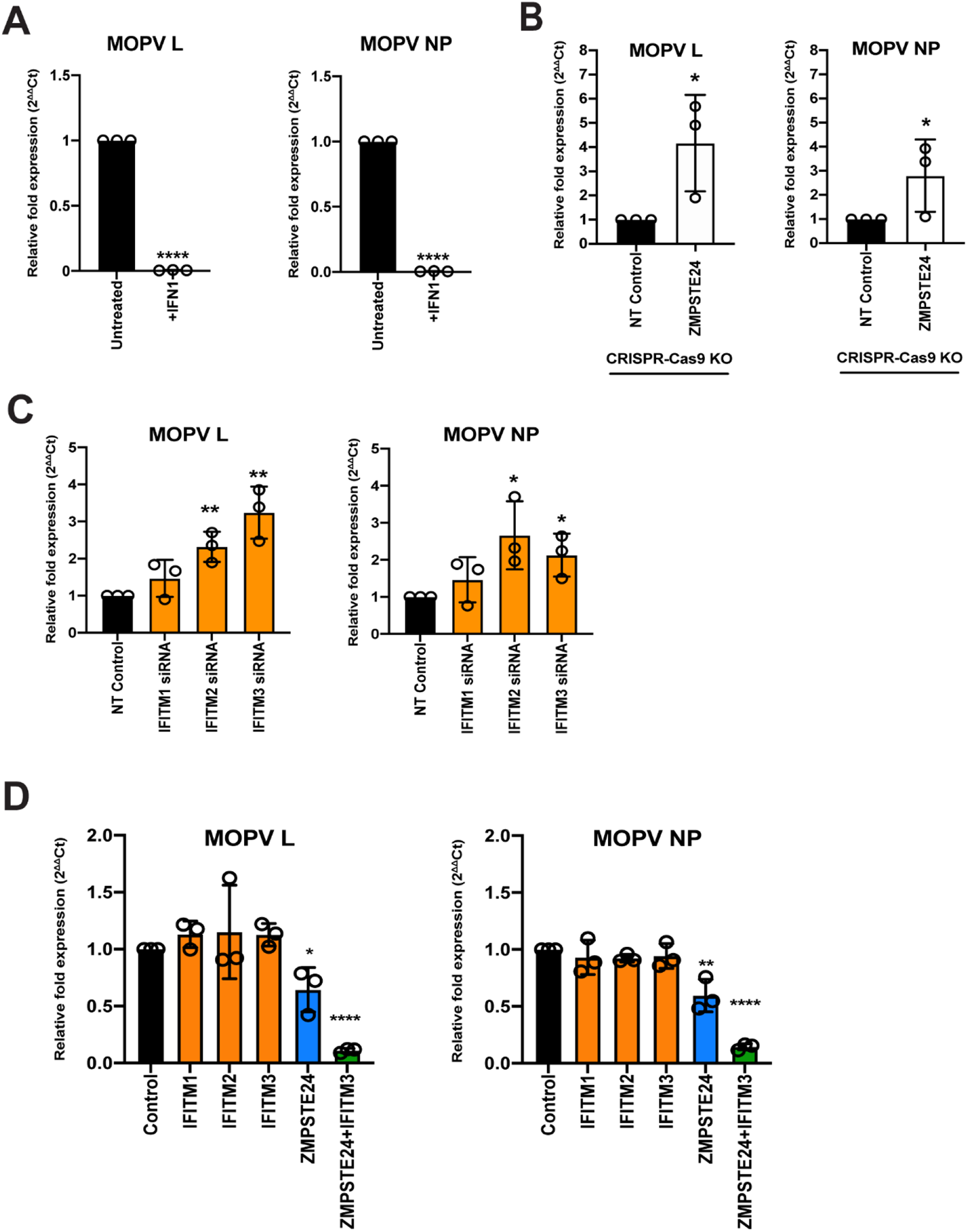
MOPV replication is sensitive to ZMPSTE24 restriction. **(A)** MOPV gene expression analysis in A549 cells incubated with 1000 U/mL type 1 IFN (IFN1) for 4 h prior to, and during infection with MOPV. Unpaired t-test ****p<0.0001. **(B)** Gene expression in A549 cells with stable CRISPR-Cas9 knockout (KO) of ZMPSTE24 infected with MOPV. Unpaired t-test *p<0.05. **(C)** MOPV gene expression in A549 cells stably expressing siRNA-mediated knockdown of IFITMs. Unpaired t-test ** p<0.01, *p<0.05. **(D)** Gene expression analysis in A549 cells stably expressing IFITMs 1, 2 or 3, ZMPSTE24 or both ZMPSTE24 and IFITM3 and infected with MOPV. All cells were infected with MOPV at MOI 0.01 for 72 h. Unpaired t-test ****p<0.0001, ** p<0.01, *p<0.05. All samples were collected by RNA extraction and cDNA synthesis before analysing by RT-qPCR with primers for MOPV L and NP genes. Data was analysed by the 2^ΔΔ^Ct method with primers for reference genes β-actin and GAPDH. Values are expressed as means ±SEM relative to controls of samples performed in triplicate.

To further address the effects of ZMPSTE24 and IFITM activity on live virus replication, we generated stable A549 cell lines expressing each human IFITM protein, ZMPSTE24 or expressing ZMPSTE24 in combination with IFITM3 (Fig S3, Fig 5D). We then infected these cells with MOPV with a multiplicity of infection (MOI) of 0.01, and found that there was little to no sensitivity to IFITM protein expression but significant sensitivity to ZMPSTE24. The reduction of MOPV gene expression in cells expressing ZMPSTE24 was further enhanced when IFITM3 was co-expressed (Fig 5D). Both single round arenavirus GPpp assays and multicycle replication studies with live MOPV v therefore show sensitivity to ZMPSTE24 restriction that is specifically enhanced in the presence of IFITM3 expression. These data are highly suggestive of an antiviral role of ZMPSTE24 and the IFITM cofactors in arenavirus cellular entry.

### ZMPSTE24 restriction of arenavirus infection is mediated through modulation of membrane integrity

The molecular mechanism of ZMPSTE24 and IFITM antiviral activity are not well characterised (33, 34, 49). Findings from previous studies suggest that these proteins restrict the fusion of viruses by altering the fluidity or curvature of the host and viral membranes, through indirect alteration of the lipid composition of the endosomal membrane or through the association with membranous co-factors, such as ZMPSTE24 and IFITM interactions (34, 50). It has previously been demonstrated that the antiviral effect of IFITM2 and IFITM3 on infection by susceptible viruses such as IAV, can be attenuated in the presence of the amphiphilic antifungal drug amphotericin B (AmphoB) (50, 51). AmphoB intercalates into endosomal membranes and indirectly abrogates IFITM-mediated restriction through enhancement of membrane fluidity (51). We therefore used AmphoB treatment to analyse whether membrane modulation is required for ZMPSTE24 restriction and for the cooperative antiviral activity of ZMPSTE24 and IFITM proteins. To address this, we used CRISPR-Cas9 ZMPSTE24 KO A549 cells that overexpressed either ZMPSTE24 or IFITM3 individually, or overexpressed both proteins in combination (Fig S2, Fig 6A). We infected these cells and CRISPR-Cas9 NT control cells with our panel of arenavirus GPpp namely LCMV, LASV, MOPV, LUJV, JUNV, MACV and CHAPV. Notably, AmphoB had no effect on arenavirus GPpp infection in NT controls. We observed that AmphoB produced a broadly significant reversal of ZMPSTE24-mediated restriction of arenavirus GPpp infection, rendering these cells less sensitive to ZMPSTE24 inhibition (Fig 6A). Furthermore, AmphoB limited the cooperative restriction of ZMPSTE24 and IFITM3 in comparison to untreated cells (Fig 6A). These data imply that at the early stages of arenavirus infection, the modulation of cellular membrane integrity is critical for the antiviral activity of ZMPSTE24 and the observed restriction enhancement in the presence of IFITM3. To examine this during live virus infection, we infected these cells with MOPV at MOI 0.01 for 72 hrs and quantified levels of MOPV L and NP gene expression. We found that AmphoB increased the sensitivity of MOPV to ZMSPTE24 alone and when expressed in combination with IFITM3 (Fig 6B). Similar to arenavirus GPpp infection (Fig 6A), expression of IFITM3 alone did not affect MOPV replication. Furthermore, we also observed no change in MOPV L and NP gene expression levels upon AmphoB treatment in these cells.

**Fig 6.**
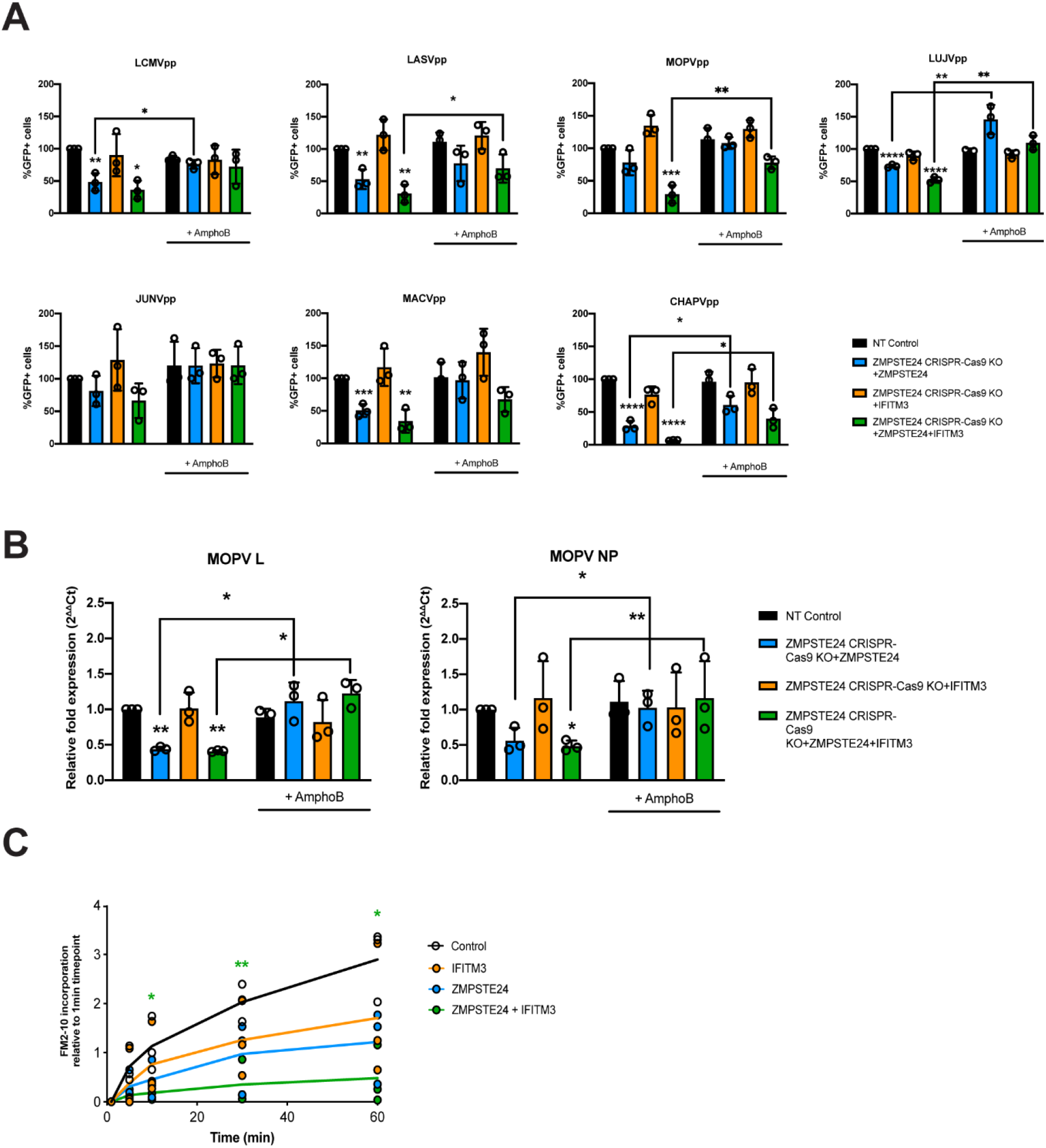
ZMPSTE24 and IFITM3 modulate membrane fluidity to restrict arenaviruses. A549 CRISPR-Cas9 stable knock out cell lines generated in Fig 4 for non-targeting (NT) control or ZMPSTE24 and then transduced for stable overexpression of ZMPSTE24-FLAG or IFITM3 or both were incubated with 1µM Amphotericin B (AmphoB) for 1 h prior to infection with **(A)** the panel of arenavirus GP-pseudoparticles (GPpp) for LCMV, LASV, MOPV, LUJV, JUNV, MACV and CHAPV for 48 h or with **(B)** MOPV (MOI 0.01) for 72 h. Infectivity of GPpp was measured by flow cytometry and calculated as percentage GFP-positive cells compared to control. Unpaired t-test ****p<0.0001, *** p<0.001, ** p<0.01, *p<0.05. MOPV L and NP gene expression was analysed by RT-qPCR and the 2^ΔΔ^Ct method with housekeeping genes β-actin and GAPDH and is displayed as relative fold change in relation to untreated NT control cells. **(C)** A549 cells stably expressing ZMPSTE24, IFITM3 or both were incubated with 200nM FM™2-10 for the indicated time points and FM™2-10 incorporation was measured as fluorescence intensity of incorporated membrane probe by flow cytometry. All data are expressed as means ±SEM from experiments performed in triplicate. Unpaired t-test ** p<0.01, *p<0.05

Structural changes in host cell membranes that can affect, for example, membrane fluidity or surface tension have implications for a wide range of biological mechanisms including cell division, endocytosis and viral fusion (50–52). Given that ZMPSTE24 and IFITMs act both independently and in synergy against a plethora of virus families, it is highly likely that they share a common mechanism that impacts on the host cellular environment. To investigate the impact of ZMPSTE24 on host membranes and indeed co-expression of the two proteins, we assessed the rate of cellular internalisation of the non-toxic membrane probe FM™2-10 in A549 cells overexpressing ZMPSTE24, IFITM3 or ZMPSTE24 and IFITM3 in combination. FM™2-10 reversibly binds to the outer leaflet of the cell membrane and upon endocytosis localises to the membrane of the endocytic vesicle (53). The fluorescence emission of FM™2-10 increases with membrane incorporation, thus we measured the changes in FM™2-10 fluorescence intensity over time by flow cytometry to determine the kinetics of membrane internalisation. In contrast to control cells, the rate and intensity of FM™2-10 probe incorporation, characteristic of membrane endocytosis, was reduced in the presence of IFITM3 or ZMPSTE24 and this reduction was significantly enhanced upon co-expression of the two restriction factors (Fig 6C). These findings strengthen the argument that ZMPSTE24 and IFITMs cause changes in membrane structure and dynamics that likely impact on the efficiency of virus fusion.

Thus, ZMPSTE24-mediated restriction activity is involved in the early stages of the innate immune response to arenavirus infection and IFITMs are able to enhance effects on membrane fluidity and thus inhibition of virus infection.

## Discussion

The entry and fusion of viruses into susceptible host cells represents a fundamental step of viral pathogenesis and is a central factor in disease outcome. Investigations into the molecular and cellular mechanisms that drive cell invasion of arenaviruses have unravelled complex details surrounding receptor switching, the regulation of virus endocytosis and the conformational rearrangements within the GP structure that induce membrane fusion during entry (27, 54). However, there is still limited knowledge regarding the range of antiviral proteins that limit the entry of arenaviruses and how the activity of these proteins may be modulated by virus-specific proteins.

In this study, we identified ZMPSTE24 as an intrinsic restriction factor against arenavirus entry and replication. The antiviral impact of ZMPSTE24 on arenavirus infection was shown by demonstrating that arenavirus GPpp infection and MOPV replication are enhanced in cells with depleted ZMPSTE24 and that ectopic expression of ZMPSTE24 caused a reduction in arenavirus GPpp and MOPV infection. Recent studies have indicated a number of enveloped viruses that traffic through the cellular endosomal compartment during entry are restricted by ZMPSTE24 (33). Our findings now expand this list to include arenaviruses. The breadth of viruses impacted by ZMPSTE24 activity suggests a universal antiviral mechanism that occurs prior to virus fusion. Our data suggests that disrupting the protease activity of ZMPSTE24 does not alter the sensitivity of arenaviruses to restriction, further implying a generalised mechanism of restriction that may involve modifying host cell membrane properties to inhibit fusion pore formation. This mechanism which impairs endosomal viral membrane fusion is likely facilitated by the IFITM proteins as co-factors of ZMPSTE24 activity.

The localisation of IFITM proteins is an important determinant of the breadth of viruses that they restrict. IFITM1 is found predominantly at the plasma membrane whilst IFITMs 2 and 3 localise to endosomal compartments (21). We confirmed and expanded previous studies that show arenaviruses are insensitive to IFITM restriction (39). However, our observations also highlight that arenavirus entry and replication is enhanced upon knock down and knockout of IFITM expression. This implies that IFITMs are involved as antiviral factors against the early stages of arenavirus infection. A recent studiy by Suddala *et al.* (39) indicated that IFITM3 restricts through a proximity-based mechanism and that LASV may escape restriction by IFITM3 by entering cells through a distinct endosomal pathway lacking IFITM3 expression. Our observations support that arenaviruses are not inherently insensitive to IFITM restriction given that IFITM depletion enhances arenavirus infection (39).

In light of our findings and given that previous studies have showed that ZMPSTE24 is required for the antiviral activity of IFITMs, we explored a possible cooperative role of ZMPSTE24 and IFITMs against arenavirus infection. Our data demonstrate that endogenous ZMPSTE24 co-localises with all three IFITM proteins when they are ectopically expressed in A549 cells. Using immunoprecipitation and complementation assays, we also showed that IFITM proteins interact with ZMPSTE24.

In our present study, we provide strong evidence for the biological significance of the ZMPSTE24—IFITM interaction demonstrating that engagement with IFITM proteins enhances the sensitivity of arenaviruses to ZMPSTE24-mediated restriction. Specifically, we show that stable ectopic expression of ZMPSTE24 with IFITM3 significantly enhanced inhibition of arenavirus entry and replication. In addition, we found that in contrast to cells singly overexpressing IFITM3, ectopic co-expression of IFITM3 with ZMPSTE24 in A549 cells led to the redistribution of IFITM3 to distinct endosomal compartments that were positive for ZMPSTE24. We therefore propose the redistribution of IFITM3 to a ZMPSTE24-positive pathway along which arenaviruses enter and fuse, induces an enhanced modification of cellular membranes that thus impairs virus fusion. Supporting this hypothesis, we provide evidence that the absence of IFITMs in A549 cells expressing ZMPSTE24 leads to a reduction in the sensitivity of arenavirus GPpp to ZMPSTE24-mediated restriction. It is therefore tempting to speculate that ZMPSTE24 is able to modulate the intracellular trafficking of its IFITM co-factors to an early endosomal localisation that increases the susceptibility of normally IFITM-resistant viruses like arenaviruses.

We aimed to address the mechanism of ZMPSTE24 restriction and of the observed cooperative activity using AmphoB treatment which disrupts IFITM function and by assessing the incorporation of a membrane-sensitive probe, FM™2-10. Pre-treatment with AmphoB rescued the entry and fusion of arenavirus GPpp and live MOPV infection in cells expressing either ZMPSTE24 alone or co-expressing ZMPSTE24 and IFITM3. These findings are consistent with the notion that ZMPSTE24 may exert its inhibitory effect by modulating the curvature and increasing the rigidity of endosomal membranes, much like the IFITM proteins. Interestingly, ZMPSTE24 decreased the rate and intensity of FM™2-10 incorporation into cellular membranes and this was further abrogated in the presence of IFITM3. Given the effects that these proteins likely have on membrane fluidity, the decreased incorporation and thus the associated reduced rate of endocytosis may be an indirect effect of changes in lipid composition and the distribution of membrane components. It also provides evidence that both ZMPSTE24 and IFITM3 exert their antiviral function by increasing membrane order and rigidity, a mechanism consistent with that proposed by previous studies on IFITM3 alone (49, 51).

In summary, our study highlights a previously unexplored restriction factor strategy that contributes to our understanding of arenavirus entry mechanisms. It provides further insight into the activities of ZMPSTE24 and IFITMs and provides the opportunity to target and augment this restriction mechanism for antiviral development. Defining the critical interaction sites between ZMPSTE24 and IFITM proteins and understanding the role of arenaviral proteins in abrogating restriction is of importance.

## Materials and methods

### Cell lines and expression constructs

Human embryonic kidney 293T (293T; ATCC), kidney epithelial Vero (Vero; ATCC), human lung adenocarcinoma epithelial (A549; ATCC) and A549 cells expressing ZMPSTE24 or the individual IFITM proteins were cultured in Dulbecco’s Modified Eagle Medium (DMEM), high glucose, GlutaMAX™ Supplement (Gibco) with 10% heat inactivated FBS (Gibco) and 200 µg/ml Gentamicin (Sigma) at 37°C, 5% CO_2_.

Expression plasmids encoding for human ZMPSTE24 with and without a C-terminal FLAG-tag or HA-tag were PCR amplified and subcloned into the pQXCIP (Clontech) backbone using flanking restriction sites *AgeI* and *BamHI*. Human IFITM1, IFITM2 and IFITM3 were cloned into the pLHCX retroviral vector (Clontech) using *XhoI* and *NotI* restriction sites. All IFITM proteins were HA-tagged by PCR-based mutagenesis using the parental pLHCX-IFITM1, 2 or 3 as templates.

Arenavirus glycoproteins for LCMV, LASV, MOPV. LUJV, JUNV, MACV and CHAPV (accession numbers M22138, M15076, M33879, FJ952384, D10072, AY624355, and EU260463, respectively) were synthesized by GeneArt (ThermoFisher) and subcloned into the pI.18 expression vector (kindly gifted by Professor Janet Daly) using *KpnI* and *XhoI* as flanking restriction sites.

A549 cells stably expressing the IFITMs 1, 2 or 3 (pLHCX) or ZMPSTE24 tagged with or without a C-terminal FLAG tag (pQXCIP) or the relevant empty vector, were generated by vesicular stomatitis virus-G (VSV-G) pseudotyped retroviral transduction. Retroviral vectors were made by transfecting 293T cells with the pCMV-Gag-Pol murine leukaemia virus (MLV) packaging construct (kindly gifted by Professor Jonathan Ball), the pLHCX or pQXCIP packaging vector of interest and pCMV VSV-G using 1mg/ml PEI®-MAX (Polysciences). Media was replaced 16 h post-transfection and viral supernatants were harvested through a 0.45 µm filter, 48 h post transfection. A549 cells were incubated for 48 h with retroviral vectors following spinoculation at 400 xg for 1 h to generate the stable cell lines. The corresponding antibiotic selection was added 48 h post-transfection. Expression of proteins was assessed by western blotting. When indicated, IFN-α (universal type 1 IFN, PBL Interferon Source) stimulation was performed using 1000 U ml^−1^ for 4 h before immunoblotting.

The NanoBiT split luciferase system was used to assess interaction of IFITM3 and ZMPSTE24. Briefly IFITM3 and ZMPSTE24 were fused to the NanoBiT large (LgBiT) or small (SmBiT) subunits of NanoLuc luciferase at the N or C terminus. The coding region of IFITM3 was amplified by PCR with added *XhoI* and *NheI* restriction sites whereas ZMPSTE24 was amplified with *BglII* and *XhoI* sites. Primers used were as follows: IFITM3 C-terminal forward (5′-ATGCATGCTAGCGCCACCATGAATCACACTGTCCAAAC-3′), IFITM3 C-terminal reverse (5′-ATGCATCTCGAGCCTCCATAGGCCTGGAAGA-3′), IFITM3 N-terminal forward (5′-ATGCATCTCGAGCGGTATGAATCACACTGTCCAA-3′), IFITM3 N-terminal reverse (5′-ATGCATGCTAGCCTATCCATAGGCCTGGA-3′), ZMPSTE24 C-terminal forward (5′-ATGCATAGATCTATGGGGATGTGGGCATCG-3′), ZMPSTE24 C-terminal reverse (5′-ATGCATCTCGAGCCGTGTTGCTTCATAGTTTTC-3′), ZMPSTE24 N-terminal forward (5′-ATGCATCTCGAGCGGTATGGGGATGTGGGCATC-3′) and ZMPSTE24 N-terminal reverse (5′-ATGCATGCTAGCTCAGTGTTGCTTCATAGT-3′). Amplified fragments were digested with corresponding restriction enzymes and ligated into the following vectors: pBiT1.1-C [TK LgBiT], pBiT2.1-C [TK SmBiT], pBiT1.1-N [TK LgBiT] and pBiT2.1-N [TK SmBiT].

### Passage and titration of Mopeia virus (MOPV)

The UVE/MOPV/UNK/MZ/Mozambique 20410 strain of MOPV was obtained from European Virus Archive and mycoplasma-free virus stocks were propagated in Vero cells in DMEM supplemented with 2% FCS. The titre of MOPV stocks was determined by focus forming assay. Vero cells were infected with serial dilutions of MOPV for 1 h at 37°C and then incubated in complete DMEM for 48 h. Infected foci were visualised using mouse monoclonal Anti-Arenavirus (OW) rGPC, Clone KL-AV-1B3 (BEI Resources, 1:200) followed by anti-mouse AlexaFluor488 secondary (Jackson ImmunoResearch; 1:1000). Virus titres were calculated as focus forming units per mL (FFU/mL) and MOI calculated for subsequent experiments.

### Infection with MOPV

A549 cells were infected for 1 h at 37°C with MOPV at an MOI 0.01. Media was then replaced and the cells incubated for a further 72 h at 37°C after which they were harvested for total RNA extraction. For interferon treatment, cells were treated with 1000 U ml^−1^ IFN-α (universal type 1 IFN, PBL Interferon Source) for 4 h before infection. Media was changed before infection with MOPV as above. To assess the effect of amphotericin B on the restriction of MOPV replication by ZMPSTE24 and the cooperative action with IFITMs, the relevant stable A549 cell lines were treated with 1 µM amphotericin B (Sigma Aldrich) for 1 h at 37°C prior to infection and following infection with MOPV.

### Generation of arenavirus GP retroviral pseudoparticles

To produce arenavirus GP retroviral pseudoparticles, encoding GFP, 293T cells were transfected with pCMV-MLV gag-pol, the pCMV-MLV GFP encoding an MLV-based transfer vector containing a CMV-GFP internal transcriptional unit (kindly gifted by Professor Jonathan Ball) and the pI.18 plasmid encoding the arenavirus GP of interest, at a ratio of 0.6:0.9:0.6 µg. Cells were transfected using 1 mg/ml PEI MAX® (Polysciences). Media was replaced 16 h post-transfection and viral supernatants were harvested through a 0.45µm filter 48 h post transfection. The viral supernatants were then titrated on A549 cells by flow cytometry.

### Entry assay using arenavirus GP pseudoparticles

Cells were infected with arenavirus GP retroviral pseudoparticles, encoding GFP at an MOI of 0.3 in complete growth media and incubated at 37°C for 48 h. Infected cells were analysed by flow cytometry. Samples were gated on live cells for 10,000 events and analysed for expression of GFP. To test the effect of IFN1 on arenavirus GP-mediated cell entry, cells were treated with IFN-α (universal type 1 IFN, PBL Interferon Source) for 4 h prior and throughout infection. To assess the effect of amphotericin B on the restriction of arenavirus entry by ZMPSTE24 and the co-operative action with IFITMs, relevant stable A549 cell lines were treated with 1 µM amphotericin B (Sigma Aldrich) for 1 h at 37°C prior to infection and following infection with arenavirus GP pseudoparticles.

### Flow cytometry

Cells are gently washed, detached and resuspended in 1% BSA in PBS and 0.1% (w/v) sodium azide. Flow cytometry analyses were performed using a BD FACSCanto II flow-cytometer (Becton Dickinson), collecting 10,000 events, and analysed using FlowJo software. Arenavirus glycoprotein pseudotyped virus vector infected cells were analysed for expression of GFP. Infected cell gates were set using uninfected control samples. FM™2-10 membrane probe incorporation was also measured by flow cytometry and cells gated on the PE channel and gates were set using non-treated control samples.

### siRNA knockdown of ZMPSTE24 and IFITMs

siRNA mediated knockdown of ZMPSTE24 and IFITM proteins was performed by transfection of A549 cells using Lipofectamine™RNAiMAX Transfection Reagent (ThermoFisher) according to the manufacturer’s instructions. Cells were transfected with 10 µM of either of the following SMARTpool siRNAs (Dharmacon): ON-TARGETplus Non-targeting Pool siRNA (D-001810-10-05) or ON-TARGETplus Human ZMPSTE24 (10269) siRNA (L-006104-00-0010) or ON-TARGETplus Human IFITM1 (8519) siRNA (L-019543-00-0005), ON-TARGETplus Human IFITM2 (10581) siRNA(L-020103-00-0005) or ON-TARGETplus Human IFITM3 (10410) siRNA (L-014116-01-0005). Knockdown was assessed by western blotting and cells used in functional assays 48 h post-transfection.

### CRISPR-Cas9 knockout of ZMPSTE24 and IFITM expression

CRISPR-Cas9 gRNA sequences to target human ZMPSTE24 (CACAACTAATGTGAACAGCC) and a non-targeting control (GGCCCTCTAGAAAAGTCTCG) were generated in pLentiCRISPR v2 and gRNA sequences to target human IFITMs 1, 2 and 3 (TTCTTCTCTCCTGTCAACAG) were generated in eSpCas9-LentiCRISPR v2 (Genscript). Viral stocks were generated in 293T cells. 293T cells were co-transfected with psPAX2 (Addgene), pMD2.G VSV-g and the eSpCas9-LentiCRISPRv2 construct targeting ZMPSTE24 or the IFITM proteins or a non-targeting control at a ratio of 0.6:0.9:0.6 using 1 mg/ml PEI^®^-MAX. Media was changed 16 h post transfection and viral stocks were harvested and filtered 48 h post transfection. A549 cells were then transduced with the pLentiCRISPR viruses at 400 x*g* for 1 h and cells cultured for a further 7 days in the presence of 1 μg/ml puromycin. The efficiency of knockout was determined by SDS-PAGE and western blot. The effects of CRISPR-Cas9 knockout of protein expression on arenavirus GP pseudoparticle entry and MOPV replication was determined by flow cytometry and by RT-qPCR assays.

### SDS-PAGE and Western blot analysis

Cellular samples were lysed in 2× reducing Laemmli buffer (Bio-Rad) at 100°C for 10 min. Samples were separated on 8–16% Mini-PROTEAN® TGX Precast gels (Bio-Rad) and transferred onto 0.2 µm nitrocellulose membrane (Bio-Rad). Membranes were blocked in 5% milk in PBS with 0.1% Tween^®^20 (PBS-T) for 30 min prior to incubation with specific primary antibodies: mouse anti-IFITM1 (Proteintech, 60074-1-Ig, 1:5000), rabbit anti-IFITM2 (Proteintech, 12769-1-AP, 1:5000), rabbit anti-IFITM3 (Proteintech, 11714-1-AP, 1:5000), rabbit anti-ZMPSTE24 antibody (Abcam ab38450, 1:1000), mouse anti-FLAG (Sigma, F1804, 1:2000), mouse anti-HA (Abcam ab18181, 1:5000), mouse anti-HSP90 (Invitrogen, MA1-10372, 1:10,000). All antibodies were diluted in 5% milk in PBS-T and incubated at 4°C overnight with gentle shaking. After washing membranes with PBS-T at RT, horseradish peroxidase-conjugated (HRP) horse anti-mouse IgG (CST, 7076S, 1:5000) and goat anti-rabbit IgG (CST, 7074S, 1:5000) secondary antibodies in 5% milk in PBS-T were added and membranes incubated for 1 h at RT with gentle shaking. Following washes in PBS-T, proteins were detected using SuperSignal™ West Pico PLUS Chemiluminescent Substrate (ThermoFisher).

### Immunoprecipitations

A549 cells were transfected with 2 µg pQXCIP ZMPSTE24-FLAG. 48 h post transfection cells were lysed on ice for 20 min in 50 mM Tris-HCL pH 7.4, 150 mM NaCl, 1% IGEPAL®CA-630 (Sigma), complete protease inhibitors (Roche). Lysed samples were centrifuged and supernatants were immunoprecipitated with 5µg/ml mouse monoclonal anti-IFITM2/3 antibody (Proteintech, 66081-1-Ig) for 1.5 h at 4°C. Protein G agarose (ThermoFisher) was equilibrated in lysis buffer before adding to supernatants and incubated with gentle rolling overnight at 4°C. Following extensive washes in lysis buffer, cell lysates and immunoprecipitates on beads were resuspended in 2× Laemmli buffer (Bio-Rad) and subjected to SDS-PAGE and western blot analysis.

### Immunofluorescence microscopy

A549 cells grown on coverslips and transiently transfected with HA-tagged IFITM or FLAG-or HA-tagged ZMPSTE24 proteins, were fixed with 4% paraformaldehyde (PFA) in PBS for 10 mins at RT. Cells were permeabilized with 0.1% Triton X-100 in PBS for 10 min at RT and stained overnight at 4°C with the appropriate primary antibodies (rabbit anti-ZMPSTE24, Abcam ab38450, 1:100 or mouse monoclonal anti-HA, Abcam ab18181, 1:500 or rabbit anti-EEA1, CST 2411S, 1:100 or rabbit anti-Rab9A, CST 5118T, 1:200, or rabbit anti-LAMP1, Invitrogen 14-1079-80, 1:500. All antibodies were diluted in 0.1% BSA, 0.01% Triton-X-100 in PBS. Following washes in PBS, coverslips were incubated for 1 h at RT with appropriate secondary antibodies conjugated to Alexa Fluor 488 or 594 (Molecular Probes, ThermoFisher 1:500) diluted in 0.1% BSA, 0.01% Triton-X-100 in PBS. Following incubation, coverslips were washed in PBS and mounted on glass slides using ProLong™ Diamond Antifade Mountant with DAPI (Molecular Probes, ThermoFisher). Images were acquired on a Zeiss LSM880 confocal laser scanning microscope or on a Leica DM5000 B widefield microscope. Z stacks were taken for all stained conditions and images were deconvolved with the Zeiss ZEN deconvolution software and analysed using ImageJ. Representative images are shown.

### NanoBiT protein interaction assay

A NanoBiT protein:protein interaction assay (Promega) was used to assess the interaction between ZMPSTE24 and IFITM3. A549 cells were transiently co-transfected with 100 ng in total of an N- or C-terminal LgBiT and SmBiT tagged construct using Lipofectamine™ 3000 Transfection Reagent (ThermoFisher). All possible combinations of the N-terminal and C-terminal-tagged split luciferase protein pairs were tested. All LgBiT constructs were co-transfected with the HaloTag-SmBiT (negative control) construct against which relative luminescence was measured. Luminescence was measured after 48 h using the Nano-Glo^®^ Live Cell Assay System (Promega) according to manufacturer’s instructions.

### RT-qPCR analysis

Nucleoprotein (NP) and RNA-dependent RNA polymerase (L) RNA levels were determined in cells infected with MOPV after 72 h of infection. Total RNA was isolated from infected cells using the QIAGEN RNeasy Plus Mini Kit (Qiagen, 74136) and cDNA was reverse transcribed using the Applied Biosystems™ High-Capacity cDNA Reverse Transcription Kit according to manufacturer’s instructions. For each qPCR reaction, 10ng of cDNA was used with the Applied Biosystems™ PowerUp™ SYBR™ Green Master Mix under the following conditions: 50°C 2 mins, 95°C 2 mins then 40 cycles of 95°C 15 secs, 55°C 15 secs, 72°C 1 min. Primers used were as follows: GAPDH forward (5′-ACATCGCTCAGACACCATG-3′); GAPDH reverse (5′-TGTAGTTGAGGTCAATGAAGGG-3′); β-actin forward (5′-CACCAACTGGGACGACAT-3′); β-actin reverse (5′-ACAGCCTGGATAGCAACG-3′); MOPV L forward (5′-ATCTCCTCATGCAGCCACAC-3′); MOPV L reverse (5′-GGACTGTTGGAGAGTTGCGA-3′); MOPV NP forward (5′-CCCTGGCATGTCAAGACCAT-3′); MOPV NP reverse (5′-CCCTGTGGAAGTTGCGATCT-3′). Primer specificity was confirmed by melt curve analysis. Relative fold expression of target genes was normalised to reference genes GAPDH and β-actin by the ^ΔΔ^Ct method.

### FM™2-10 incorporation assay

The effect of ZMPSTE24 and IFITM3 expression on the rate of internalisation of the FM™2-10 (N-(3-Triethylammoniumpropyl)-4-(4-(Diethylamino)styryl)Pyridinium Dibromide, ThermoFisher) membrane probe into A549 cells was determined by flow cytometry. The detection of FM™2-10 fluorescence intensity as a function of time was used a measure of endocytosis. A549 cells stably expressing ZMPSTE24, or IFITM3 or ZMPSTE24 and IFITM3 in combination or pQXCIP empty vector were washed and resuspended in PBS. A 2 µM stock solution of FM™2-10 was prepared in PBS before adding cells to a final concentration of 200 nM and incubating for 5, 10, 30 and 60min time points. The changes in FM™2-10 fluorescence intensity over time were detected by flow cytometry for each cell condition analysed.

### Statistical analysis

All statistical analyses were carried out using GraphPad Prism v9.0.2. Levels of significance were determined as follows: ****p<0.0001, *** p<0.001, ** p<0.01, *p<0.05. Data was subjected to independent sample t-tests.

## Acknowledgements

We are grateful to Professor Janet Daly for support and helpful advice. We thank Professor Jonathan Ball for helpful discussions. We are also grateful to Dr Thomas Strecker for advice and helpful suggestions. We thank Sam Stafford at Leeds Beckett University for his help with deconvolution of microscopy images. We also thank the members of the School of Veterinary Medicine and Science, University of Nottingham for their collegiate nature during a challenging year.

This research was supported by a Wellcome Trust Seed Award in Science 217414/Z/19/Z and Nottingham Research Fellowship to TLF.

All experiments were planned and performed by RJS and TLF. Confocal microscopy was carried out by the University of Nottingham, School of Life Sciences Imaging Unit (SLIM). RJS and TLF analysed the data. RJS and TLF wrote the manuscript. All authors edited the manuscript and provided comments.

## References

1. McLay L, Liang Y, Ly H. 2014. Comparative analysis of disease pathogenesis and molecular mechanisms of New World and Old World arenavirus infections. J Gen Virol 95:1–15.

2. Wolff H, Lange JV, Webb PA. 1978. Interrelationships among arenaviruses measured by indirect immunofluorescence. Intervirology 9:344–50.

3. Bowen MD, Peters CJ, Nichol ST. 1996. The Phylogeny of New World (Tacaribe Complex) Arenaviruses. Virology 219:285–290.

4. Clegg JC. 2002. Molecular phylogeny of the arenaviruses. Curr Top Microbiol Immunol 262:1–24.

5. Briese T, Paweska JT, McMullan LK, Hutchison SK, Street C, Palacios G, Khristova ML, Weyer J, Swanepoel R, Egholm M, Nichol ST, Lipkin WI. 2009. Genetic detection and characterization of Lujo virus, a new hemorrhagic fever-associated arenavirus from southern Africa. PLoS Pathog 5:e1000455.

6. Ilori EA, Furuse Y, Ipadeola OB, Dan-Nwafor CC, Abubakar A, Womi-Eteng OE, Ogbaini-Emovon E, Okogbenin S, Unigwe U, Ogah E, Ayodeji O, Abejegah C, Liasu AA, Musa EO, Woldetsadik SF, Lasuba CLP, Alemu W, Ihekweazu C. 2019. Epidemiologic and Clinical Features of Lassa Fever Outbreak in Nigeria, January 1-May 6, 2018. Emerg Infect Dis 25:1066–1074.

7. Kafetzopoulou LE, Pullan ST, Lemey P, Suchard MA, Ehichioya DU, Pahlmann M, Thielebein A, Hinzmann J, Oestereich L, Wozniak DM, Efthymiadis K, Schachten D, Koenig F, Matjeschk J, Lorenzen S, Lumley S, Ighodalo Y, Adomeh DI, Olokor T, Omomoh E, Omiunu R, Agbukor J, Ebo B, Aiyepada J, Ebhodaghe P, Osiemi B, Ehikhametalor S, Akhilomen P, Airende M, Esumeh R, Muoebonam E, Giwa R, Ekanem A, Igenegbale G, Odigie G, Okonofua G, Enigbe R, Oyakhilome J, Yerumoh EO, Odia I, Aire C, Okonofua M, Atafo R, Tobin E, Asogun D, Akpede N, Okokhere PO, Rafiu MO, Iraoyah KO, Iruolagbe CO, et al. 2019. Metagenomic sequencing at the epicenter of the Nigeria 2018 Lassa fever outbreak. Science 363:74–77.

8. Oloniniyi OK, Unigwe US, Okada S, Kimura M, Koyano S, Miyazaki Y, Iroezindu MO, Ajayi NA, Chukwubike CM, Chika-Igwenyi NM, Ndu AC, Nwidi DU, Abe H, Urata S, Kurosaki Y, Yasuda J. 2018. Genetic characterization of Lassa virus strains isolated from 2012 to 2016 in southeastern Nigeria. PLoS Negl Trop Dis 12:e0006971.

9. Siddle KJ, Eromon P, Barnes KG, Mehta S, Oguzie JU, Odia I, Schaffner SF, Winnicki SM, Shah RR, Qu J, Wohl S, Brehio P, Iruolagbe C, Aiyepada J, Uyigue E, Akhilomen P, Okonofua G, Ye S, Kayode T, Ajogbasile F, Uwanibe J, Gaye A, Momoh M, Chak B, Kotliar D, Carter A, Gladden-Young A, Freije CA, Omoregie O, Osiemi B, Muoebonam EB, Airende M, Enigbe R, Ebo B, Nosamiefan I, Oluniyi P, Nekoui M, Ogbaini-Emovon E, Garry RF, Andersen KG, Park DJ, Yozwiak NL, Akpede G, Ihekweazu C, Tomori O, Okogbenin S, Folarin OA, Okokhere PO, MacInnis BL, Sabeti PC, et al. 2018. Genomic Analysis of Lassa Virus during an Increase in Cases in Nigeria in 2018. N Engl J Med 379:1745–1753.

10. Asogun DA, Günther S, Akpede GO, Ihekweazu C, Zumla A. 2019. Lassa Fever: Epidemiology, Clinical Features, Diagnosis, Management and Prevention. Infect Dis Clin North Am 33:933–951.

11. Control NCfD. 2021. Lassa fever Situation Report Epi Week 10: 8 – 14 March 2021, p5, https://ncdc.gov.ng/diseases/sitreps/?cat=5&name=An%20update%20of%20Lassa%20fever%20outbreak%20in%20Nigeria.

12. Kofman A, Choi M, Rollin P. 2019. Lassa Fever in Travelers from West Africa, 1969–2016. Emerging Infectious Disease journal 25:236.

13. Overbosch F, de Boer M, Veldkamp KE, Ellerbroek P, Bleeker-Rovers CP, Goorhuis B, van Vugt M, van der Eijk A, Leenstra T, Khargi M, Ros J, Brandwagt D, Haverkate M, Swaan C, Reusken C, Timen A, Koopmans M, van Dissel J. 2020. Public health response to two imported, epidemiologically related cases of Lassa fever in the Netherlands (ex Sierra Leone), November 2019. Euro Surveill 25.

14. Whitmer SLM, Strecker T, Cadar D, Dienes HP, Faber K, Patel K, Brown SM, Davis WG, Klena JD, Rollin PE, Schmidt-Chanasit J, Fichet-Calvet E, Noack B, Emmerich P, Rieger T, Wolff S, Fehling SK, Eickmann M, Mengel JP, Schultze T, Hain T, Ampofo W, Bonney K, Aryeequaye JND, Ribner B, Varkey JB, Mehta AK, Lyon GM, 3rd, Kann G, De Leuw P, Schuettfort G, Stephan C, Wieland U, Fries JWU, Kochanek M, Kraft CS, Wolf T, Nichol ST, Becker S, Ströher U, Günther S. 2018. New Lineage of Lassa Virus, Togo, 2016. Emerg Infect Dis 24:599–602.

15. Wolff S, Schultze T, Fehling SK, Mengel JP, Kann G, Wolf T, Eickmann M, Becker S, Hain T, Strecker T. 2016. Genome Sequence of Lassa Virus Isolated from the First Domestically Acquired Case in Germany. Genome Announc 4.

16. Carnec X, Mateo M, Page A, Reynard S, Hortion J, Picard C, Yekwa E, Barrot L, Barron S, Vallve A, Raoul H, Carbonnelle C, Ferron F, Baize S. 2018. A Vaccine Platform against Arenaviruses Based on a Recombinant Hyperattenuated Mopeia Virus Expressing Heterologous Glycoproteins. J Virol 92.

17. Kiley MP, Lange JV, Johnson KM. 1979. Protection of rhesus monkeys from Lassa virus by immunisation with closely related Arenavirus. Lancet 2:738.

18. Walker DH, Johnson KM, Lange JV, Gardner JJ, Kiley MP, McCormick JB. 1982. Experimental infection of rhesus monkeys with Lassa virus and a closely related arenavirus, Mozambique virus. J Infect Dis 146:360–8.

19. Mazzola LT Kelly-Cirino C. 2019. Diagnostics for Lassa fever virus: a genetically diverse pathogen found in low-resource settings. BMJ Global Health 4:e001116.

20. Bieniasz PD. 2004. Intrinsic immunity: a front-line defense against viral attack. Nature Immunology 5:1109–1115.

21. Foster TL, Wilson H, Iyer SS, Coss K, Doores K, Smith S, Kellam P, Finzi A, Borrow P, Hahn BH, Neil SJD. 2016. Resistance of Transmitted Founder HIV-1 to IFITM-Mediated Restriction. Cell Host Microbe 20:429–442.

22. Shi G, Kenney AD, Kudryashova E, Zani A, Zhang L, Lai KK, Hall-Stoodley L, Robinson RT, Kudryashov DS, Compton AA, Yount JS. 2021. Opposing activities of IFITM proteins in SARS-CoV-2 infection. Embo j 40:e106501.

23. Shi G, Schwartz O, Compton AA. 2017. More than meets the I: the diverse antiviral and cellular functions of interferon-induced transmembrane proteins. Retrovirology 14:53.

24. Winstone H, Lista MJ, Reid AC, Bouton C, Pickering S, Galao RP, Kerridge C, Doores KJ, Swanson C, Neil S. 2021. The polybasic cleavage site in the SARS-CoV-2 spike modulates viral sensitivity to Type I interferon and IFITM2. J Virol doi:10.1128/jvi.02422-20.

25. Abraham J, Kwong JA, Albariño CG, Lu JG, Radoshitzky SR, Salazar-Bravo J, Farzan M, Spiropoulou CF, Choe H. 2009. Host-Species Transferrin Receptor 1 Orthologs Are Cellular Receptors for Nonpathogenic New World Clade B Arenaviruses. PLOS Pathogens 5:e1000358.

26. Jae LT, Raaben M, Herbert AS, Kuehne AI, Wirchnianski AS, Soh TK, Stubbs SH, Janssen H, Damme M, Saftig P, Whelan SP, Dye JM, Brummelkamp TR. 2014. Virus entry. Lassa virus entry requires a trigger-induced receptor switch. Science 344:1506–10.

27. Li S, Sun Z, Pryce R, Parsy M-L, Fehling SK, Schlie K, Siebert CA, Garten W, Bowden TA, Strecker T, Huiskonen JT. 2016. Acidic pH-Induced Conformations and LAMP1 Binding of the Lassa Virus Glycoprotein Spike. PLoS pathogens 12:e1005418–e1005418.

28. Raaben M, Jae LT, Herbert AS, Kuehne AI, Stubbs SH, Chou YY, Blomen VA, Kirchhausen T, Dye JM, Brummelkamp TR, Whelan SP. 2017. NRP2 and CD63 Are Host Factors for Lujo Virus Cell Entry. Cell Host Microbe 22:688–696.e5.

29. Bulow U, Govindan R, Munro JB. 2020. Acidic pH Triggers Lipid Mixing Mediated by Lassa Virus GP. Viruses 12.

30. Di Simone C, Buchmeier MJ. 1995. Kinetics and pH dependence of acid-induced structural changes in the lymphocytic choriomeningitis virus glycoprotein complex. Virology 209:3–9.

31. Chemudupati M, Kenney AD, Bonifati S, Zani A, McMichael TM, Wu L, Yount JS. 2019. From APOBEC to ZAP: Diverse mechanisms used by cellular restriction factors to inhibit virus infections. Biochimica et Biophysica Acta (BBA) – Molecular Cell Research 1866:382–394.

32. Stott RJ, Strecker T, Foster TL. 2020. Distinct Molecular Mechanisms of Host Immune Response Modulation by Arenavirus NP and Z Proteins. Viruses 12.

33. Fu B, Wang L, Li S, Dorf ME. 2017. ZMPSTE24 defends against influenza and other pathogenic viruses. Journal of Experimental Medicine 214:919–929.

34. Li S, Fu B, Wang L, Dorf ME. 2017. ZMPSTE24 Is Downstream Effector of Interferon-Induced Transmembrane Antiviral Activity. DNA Cell Biol 36:513–517.

35. Brass AL, Huang IC, Benita Y, John SP, Krishnan MN, Feeley EM, Ryan BJ, Weyer JL, van der Weyden L, Fikrig E, Adams DJ, Xavier RJ, Farzan M, Elledge SJ. 2009. The IFITM Proteins Mediate Cellular Resistance to Influenza A H1N1 Virus, West Nile Virus, and Dengue Virus. Cell 139:1243–1254.

36. Desai TM, Marin M, Chin CR, Savidis G, Brass AL, Melikyan GB. 2014. IFITM3 restricts influenza A virus entry by blocking the formation of fusion pores following virus-endosome hemifusion. PLoS Pathog 10:e1004048.

37. Everitt AR, Clare S, McDonald JU, Kane L, Harcourt K, Ahras M, Lall A, Hale C, Rodgers A, Young DB, Haque A, Billker O, Tregoning JS, Dougan G, Kellam P. 2013. Defining the Range of Pathogens Susceptible to Ifitm3 Restriction Using a Knockout Mouse Model. PLOS ONE 8:e80723.

38. Mudhasani R, Tran JP, Retterer C, Radoshitzky SR, Kota KP, Altamura LA, Smith JM, Packard BZ, Kuhn JH, Costantino J, Garrison AR, Schmaljohn CS, Huang IC, Farzan M, Bavari S. 2013. IFITM-2 and IFITM-3 but not IFITM-1 restrict Rift Valley fever virus. J Virol 87:8451–64.

39. Suddala KC, Lee CC, Meraner P, Marin M, Markosyan RM, Desai TM, Cohen FS, Brass AL, Melikyan GB. 2019. Interferon-induced transmembrane protein 3 blocks fusion of sensitive but not resistant viruses by partitioning into virus-carrying endosomes. PLoS Pathog 15:e1007532.

40. Weston S, Czieso S, White IJ, Smith SE, Kellam P, Marsh M. 2014. A Membrane Topology Model for Human Interferon Inducible Transmembrane Protein 1. PLOS ONE 9:e104341.

41. John SP, Chin CR, Perreira JM, Feeley EM, Aker AM, Savidis G, Smith SE, Elia AEH, Everitt AR, Vora M, Pertel T, Elledge SJ, Kellam P, Brass AL. 2013. The CD225 domain of IFITM3 is required for both IFITM protein association and inhibition of influenza A virus and dengue virus replication. Journal of virology 87:7837–7852.

42. Barrowman J, Michaelis S. 2009. ZMPSTE24, an integral membrane zinc metalloprotease with a connection to progeroid disorders. Biol Chem 390:761–73.

43. York J, Nunberg JH. 2006. Role of the stable signal peptide of Junín arenavirus envelope glycoprotein in pH-dependent membrane fusion. J Virol 80:7775–80.

44. Radoshitzky SR, Abraham J, Spiropoulou CF, Kuhn JH, Nguyen D, Li W, Nagel J, Schmidt PJ, Nunberg JH, Andrews NC, Farzan M, Choe H. 2007. Transferrin receptor 1 is a cellular receptor for New World haemorrhagic fever arenaviruses. Nature 446:92–6.

45. Rojek JM, Sanchez AB, Nguyen NT, de la Torre JC, Kunz S. 2008. Different mechanisms of cell entry by human-pathogenic Old World and New World arenaviruses. J Virol 82:7677–87.

46. Chen D, Hou Z, Jiang D, Zheng M, Li G, Zhang Y, Li R, Lin H, Chang J, Zeng H, Guo J-T, Zhao X. 2019. GILT restricts the cellular entry mediated by the envelope glycoproteins of SARS-CoV, Ebola virus and Lassa fever virus. Emerging Microbes & Infections 8:1511–1523.

47. Radoshitzky SR, Dong L, Chi X, Clester JC, Retterer C, Spurgers K, Kuhn JH, Sandwick S, Ruthel G, Kota K, Boltz D, Warren T, Kranzusch PJ, Whelan SP, Bavari S. 2010. Infectious Lassa virus, but not filoviruses, is restricted by BST-2/tetherin. J Virol 84:10569–80.

48. Huang IC, Bailey CC, Weyer JL, Radoshitzky SR, Becker MM, Chiang JJ, Brass AL, Ahmed AA, Chi X, Dong L, Longobardi LE, Boltz D, Kuhn JH, Elledge SJ, Bavari S, Denison MR, Choe H, Farzan M. 2011. Distinct Patterns of IFITM-Mediated Restriction of Filoviruses, SARS Coronavirus, and Influenza A Virus. PLOS Pathogens 7:e1001258.

49. Li K, Markosyan RM, Zheng YM, Golfetto O, Bungart B, Li M, Ding S, He Y, Liang C, Lee JC, Gratton E, Cohen FS, Liu SL. 2013. IFITM proteins restrict viral membrane hemifusion. PLoS Pathog 9:e1003124.

50. Wrensch F, Ligat G, Heydmann L, Schuster C, Zeisel MB, Pessaux P, Habersetzer F, King BJ, Tarr AW, Ball JK, Winkler M, Pöhlmann S, Keck ZY, Foung SKH, Baumert TF. 2019. Interferon-Induced Transmembrane Proteins Mediate Viral Evasion in Acute and Chronic Hepatitis C Virus Infection. Hepatology 70:1506–1520.

51. Lin TY, Chin CR, Everitt AR, Clare S, Perreira JM, Savidis G, Aker AM, John SP, Sarlah D, Carreira EM, Elledge SJ, Kellam P, Brass AL. 2013. Amphotericin B increases influenza A virus infection by preventing IFITM3-mediated restriction. Cell Rep 5:895–908.

52. Torriani G, Trofimenko E, Mayor J, Fedeli C, Moreno H, Michel S, Heulot M, Chevalier N, Zimmer G, Shrestha N, Plattet P, Engler O, Rothenberger S, Widmann C, Kunz S. 2019. Identification of Clotrimazole Derivatives as Specific Inhibitors of Arenavirus Fusion. J Virol 93.

53. Richards DA, Guatimosim C, Betz WJ. 2000. Two endocytic recycling routes selectively fill two vesicle pools in frog motor nerve terminals. Neuron 27:551–9.

54. Hulseberg CE, Fénéant L, Szymańska KM, White JM. 2018. Lamp1 Increases the Efficienicy of Lassa Virus Infection by Promoting Fusion in Less Acidic Endosomal Compartments. mBio 9.

