## Supplemental Material for "Inhibition of arenavirus entry and replication by the cell-intrinsic restriction factor ZMPSTE24 is enhanced by IFITM antiviral activity"

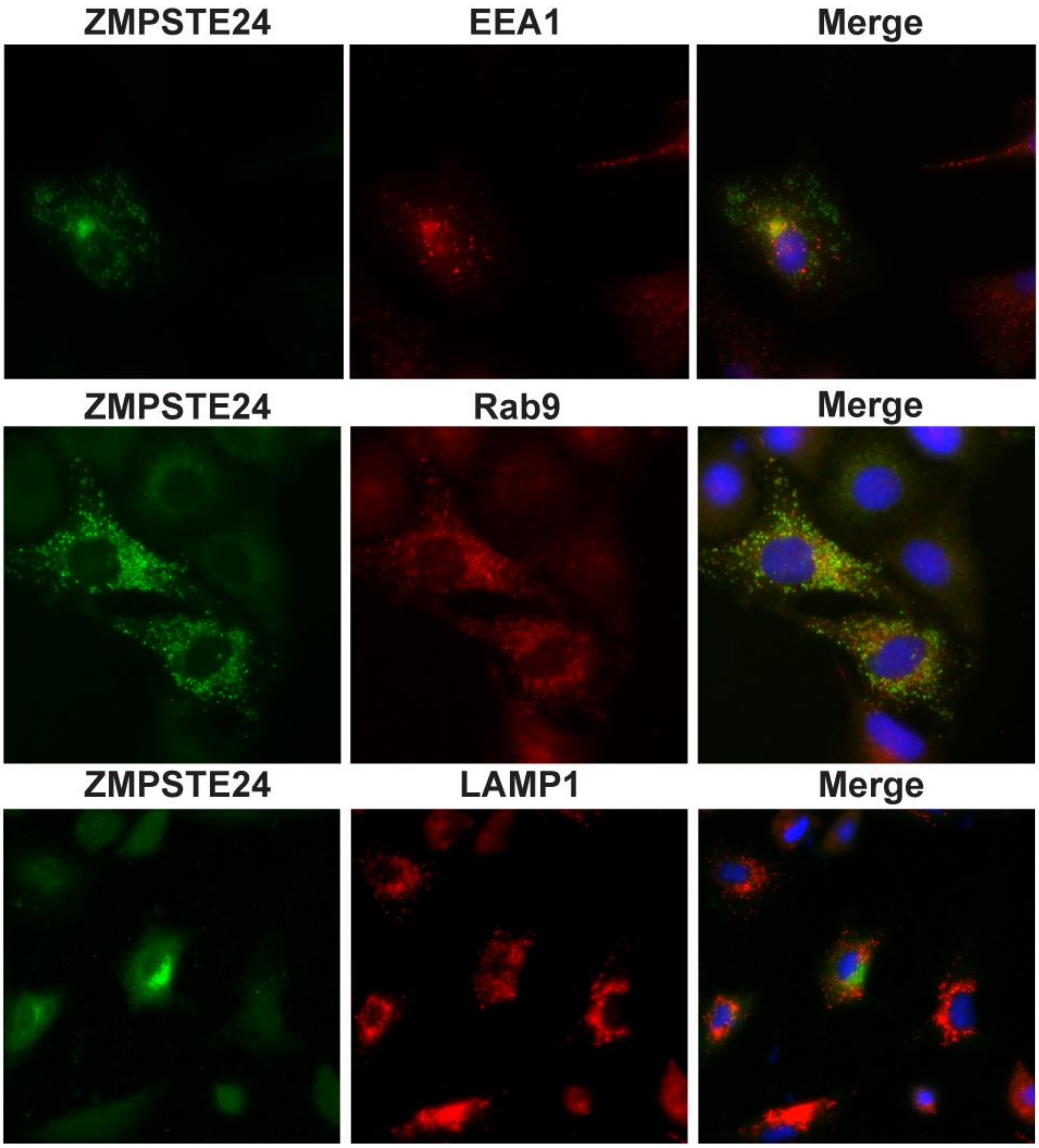


**Fig S1**

A549 cells were transiently transfected with ZMPSTE24-HA and fixed, permeabilised and immunostained with anti-HA (green) in combination with anti-EEA1, anti-Rab9 or anti-LAMP1 (red) and counterstained with DAPI (blue).

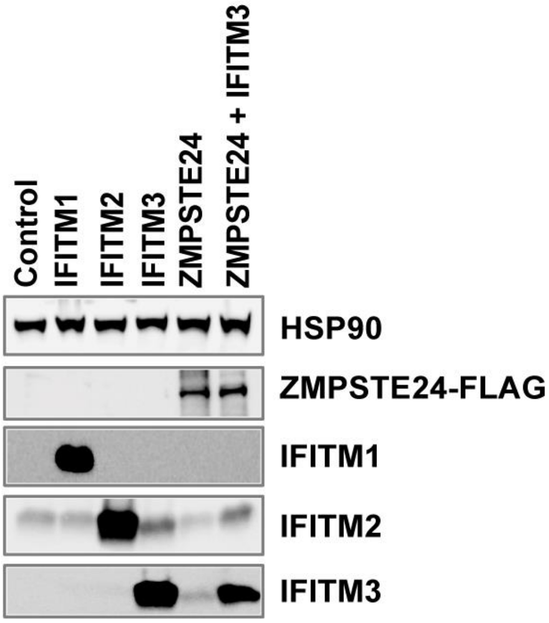


**Fig S2**

Western blot analysis of ectopic stable expression of IFITMs 1, 2 and 3 and ZMPSTE24-FLAG expression in A549 cells. Lysates were separated by SDS-PAGE and western blot then labelled using anti-FLAG, anti-IFITM1, anti-IFITM2, anti-IFITM3 or anti-HSP90 as a loading control.

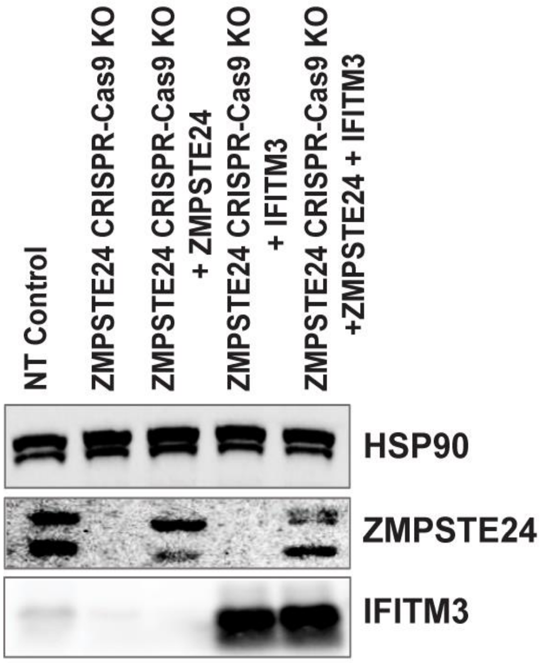


**Fig S3**

The efficiency of CRISPR-Cas9 KO of endogenous ZMPSTE24 was assessed by western blotting along with ectopic expression of ZMPSTE24-FLAG, IFITM3 or the combination of the two proteins achieved by retroviral transduction. Cell lysates were analysed by SDS-PAGE and western blotting with anti-ZMPSTE24, anti-IFITM3 or anti-HSP90 which acted as a loading control.
